# Complex-I Preserves Mitochondrial Polarization during Infection of Human Macrophages by Secretion-competent Bacteria

**DOI:** 10.1101/2025.09.04.674018

**Authors:** Francisco-Javier Garcia-Rodriguez, Paula Martinez-Oca, Carmen Buchrieser, Pedro Escoll

## Abstract

Intracellular bacteria remodel host bioenergetics and modulate mitochondrial membrane potential (Δψm). However, how individual electron-transport chain (ETC) components sustain Δψm during infection of primary human macrophages remains unclear. Here we combine extracellular flux analysis with single-cell live imaging to understand how the ETC functions in human monocyte-derived macrophages (hMDMs) during infection with (*Legionella pneumophila* (*Lp*) or *Salmonella enterica* serovar Typhimurium (*S*.Tm). At 5 h post-infection, the *Lp* type IV secretion system (T4SS) and the *S*.Tm SPI-1 T3SS were required for the early drop of the oxygen consumption rate. Despite reduced respiration, the Δψm was preserved in all infection conditions and pathogen-specific strategies to maintain the Δψm were revealed. While *Lp* infection modulates the F_O_F_1_-ATPase to function in the reverse mode (hydrolase) with the adenine-nucleotide translocator (ANT) remaining in forward mode, *S*.Tm does not reverse the F_O_F_1_-ATPase during infection. Systematic inhibition of ETC complexes established that Complex I is uniquely required to maintain the Δψm during infection with virulent bacteria but not with secretion-deficient mutant strains. Complex II is required in all infection conditions but its inhibition had a minimal effect in non-infected cells, indicating infection-driven participation of this complex in the electron flow in the ETC coupled with the preservation of the Δψm. Complexes III and IV were essential in infected and non-infected cells. Together, our results identify a Complex I-driven maintenance of the Δψm, establishing Complex I as a bioenergetic checkpoint that distinguishes virulent from secretion-deficient intracellular bacteria. Furthermore we reveal that divergent strategies are employed by *Lp* and *S*.Tm to preserve macrophage mitochondrial polarization early during infection.

## Introduction

Intracellular bacterial pathogens reprogram the metabolism of the host cells they infect to acquire nutrients to replicate inside cells ^1–3^. Macrophages, which are sentinel innate immune cells distributed across tissues, are frequent targets of intracellular bacteria and their metabolic manipulations ^4–6^. Among intracellular bacteria that infect macrophages and cause human disease, *Legionella pneumophila* (*Lp*), the causative agent of legionellosis, and *Salmonella enterica* serovar Typhimurium (*S*.Tm), the causative agent of salmonellosis, have evolved mechanisms to manipulate the host cell metabolism in their advantage ^4,7,8^. Mitochondria are at the center of these metabolic host-pathogen interactions, as these organelles coordinate energy production and other important processes, such as shaping inflammatory outputs and influencing cell death decisions ^9,10,1^. Consequently, mitochondria are important targets for intracellular bacteria^9,11–13^.

*Lp* replicates within lung macrophages and translocates over 330 bacterial effectors via its type IV secretion system (T4SS) named Dot/Icm, into its host cell to remodel a multitude of host cell processes and to establish a *Legionella*-containing vacuole (LCV) ^14^. In contrast, *S*.Tm relies on two T3SS, SPI-1 and SPI-2, to invade cells and to build and maintain the *Salmonella*-containing vacuole (SCV) in which it survives and replicates ^15^. Both, *Lp* and *S*.Tm manipulate mitochondrial functions by secreting bacterial effectors that target mitochondria ^7,11,16^.

Energy production in mitochondria relies on the generation of ATP via oxidative phosphorylation (OxPhos). At a mechanistic level, OxPhos couples electron transfer through the respiratory chain (Complex I to Complex IV) to proton pumping across the inner mitochondrial membrane, generating the proton-motive force that drives ATP synthesis by the mitochondrial F_O_F_1_-ATPase (Complex V). Several intracellular bacteria target these processes during infection ^17^, prompting the question of how mitochondrial electron transport chain (ETC) components are rewired in infected macrophages.

Previous work established that *Lp* rapidly reduces OxPhos in a T4SS-dependent manner and induces a “Warburg-like” metabolic state ^18^. This is partly achieved by the effector MitF, which induces the recruitment of host GTPase DRP-1 to mitochondria causing mitochondrial fragmentation and reducing respiration during the first hours post-infection, and before bacterial replication begins ^18^. Despite reduced respiration, *Lp* uses the T4SS effector LpSpl to maintain the Δψm by forcing the mitochondrial F_O_F_1_-ATPase to operate in the ATP-hydrolyzing, reverse mode. This delays host cell death and preserves the replication niche, the infected macrophage cell ^19^. In *S*.Tm the T3SS effector SopE2 is involved in the metabolic rewiring of infected macrophages leading to reduced mitochondrial respiration during infection ^20^. The T3SS effector SipA avoids DRP-1 recruitment to mitochondria, which blocks mitochondrial fragmentation and preserves Δψm of the infected cell ^16^. Thus, *Lp* uses one effector (MitF) to reduce mitochondrial respiration via mitochondrial fragmentation and another effector (LpSpl) to avoid the consequences of halting mitochondrial respiration (cell death) to help maintaining the Δψm. In contrast *S*.Tm uses one effector (SopE2) to reduce mitochondrial respiration and uses another effector (SipA) to maintain the Δψm via blocking mitochondrial fragmentation. This highlights a convergence of both pathogens: reducing mitochondrial respiration while preserving the Δψm, which delays cell death. However, this shared strategy is accomplished by different mechanisms in each bacterium. It is not known how each specific ETC complex in the mitochondrial OxPhos machinery (Complexes I-IV and the F_O_F_1_-ATPase) contributes to the maintenance of the Δψm during *Lp* and *S*.Tm infection of macrophages.

Here we address this question by systematically investigating the contribution of each ETC complex during *Lp* and *S*.Tm infection of human primary macrophages. Building on the evidence that *Lp* and *S*.Tm reduce oxygen consumption while they preserve the Δψm in a secretion-dependent manner, we have combined extracellular flux analyses, single-cell measurements of the Δψm and mROS, and targeted perturbations of ETC complexes to uncover the specific contribution of each mitochondrial ETC complex and each bacterial secretion system in the preservation of mitochondrial polarization during infection of macrophages by the intracellular bacteria *Lp* and *S*.Tm.

## Results and Discussion

### Only secretion-competent Legionella pneumophila and Salmonella enterica serovar Typhimurium reduce mitochondrial respiration in human macrophages

To assess how mitochondrial respiration is affected during *Lp* or *S*.Tm infection of human macrophages and if it depends on bacterial secretion systems, we infected hMDMs with *Lp* wild type (WT) or its isogenic T4SS-deficient mutant (Δ*dotA*), and with *S*.Tm WT or its isogenic T3SS-SPI-1-(Δ*prgH*), T3SS-SPI-2-(Δ*ssaV*) or T3SS-SPI-1/SPI-2-deficient double mutant (Δ*prgH*/Δ*ssaV*), and measured the oxygen consumption rate (OCR) using the Seahorse technology at 5 h post-infection (hpi), an early time point where bacteria have not replicated yet ^18,21^. Basal OCR dropped markedly in hMDMs infected with *Lp*-WT compared to non-infected controls. In contrast the Δ*dotA* mutant did not reduce the OCR and showed higher values than non-infected cells (**Figure 1A**), as previously shown ^18^, indicating that the early drop in Ox-Phos requires T4SS effectors. The basal OCR was also reduced by *S*.Tm-WT (**Figure 1B**). Loss of the SPI-1 T3SS (Δ*prgH*) or of both SPI-1 and SPI-2 (Δ*prgH*/Δ*ssaV*) abolished this reduction to values similar to non-infected cells, whereas Δ*ssaV* (SPI-2 T3SS) still showed a low OCR similar to that of the WT. Hence, the OCR reduction in WT-infected hMDMs is SPI-1–dependent, but the SPI-2 is dispensable in this assay. This aligns with published results showing that *S*.Tm reprograms the metabolism of murine macrophages towards a Warburg-like metabolic state through the SPI-1 effector SopE2, with decreased OxPhos measured in infected macrophages ^20^. In addition to a lower basal OCR at 5 hpi, *Lp*-WT- and *S*.Tm-WT-infected cells also showed a reduced FCCP response relative to their secretion-defective controls (Δ*dotA*, Δ*prgH*/Δ*ssaV*). As FCCP dissipates Δψm mimicking a physiological energy demand by stimulating the ETC to operate at maximum capacity, our results indicated a reduced maximal respiration and spare respiratory capacity during infection with WT strains. Oligomycin, an inhibitor of the F_O_F_1_-ATPase, lowered the OCR to similar minima across conditions, and addition of Rotenone + Antimycin A, inhibitors of Complex I and Complex III, respectively, collapsed OCR to non-mitochondrial levels, confirming assay specificity (**Figure 1C** and **1D**). Collectively, our results show that secretion-competent strains of two different intracellular bacteria converge on an early, robust reduction of mitochondrial respiration in primary human macrophages, T4SS for *Legionella* and predominantly SPI-1 T3SS for *Salmonella*.

**Figure 1.**
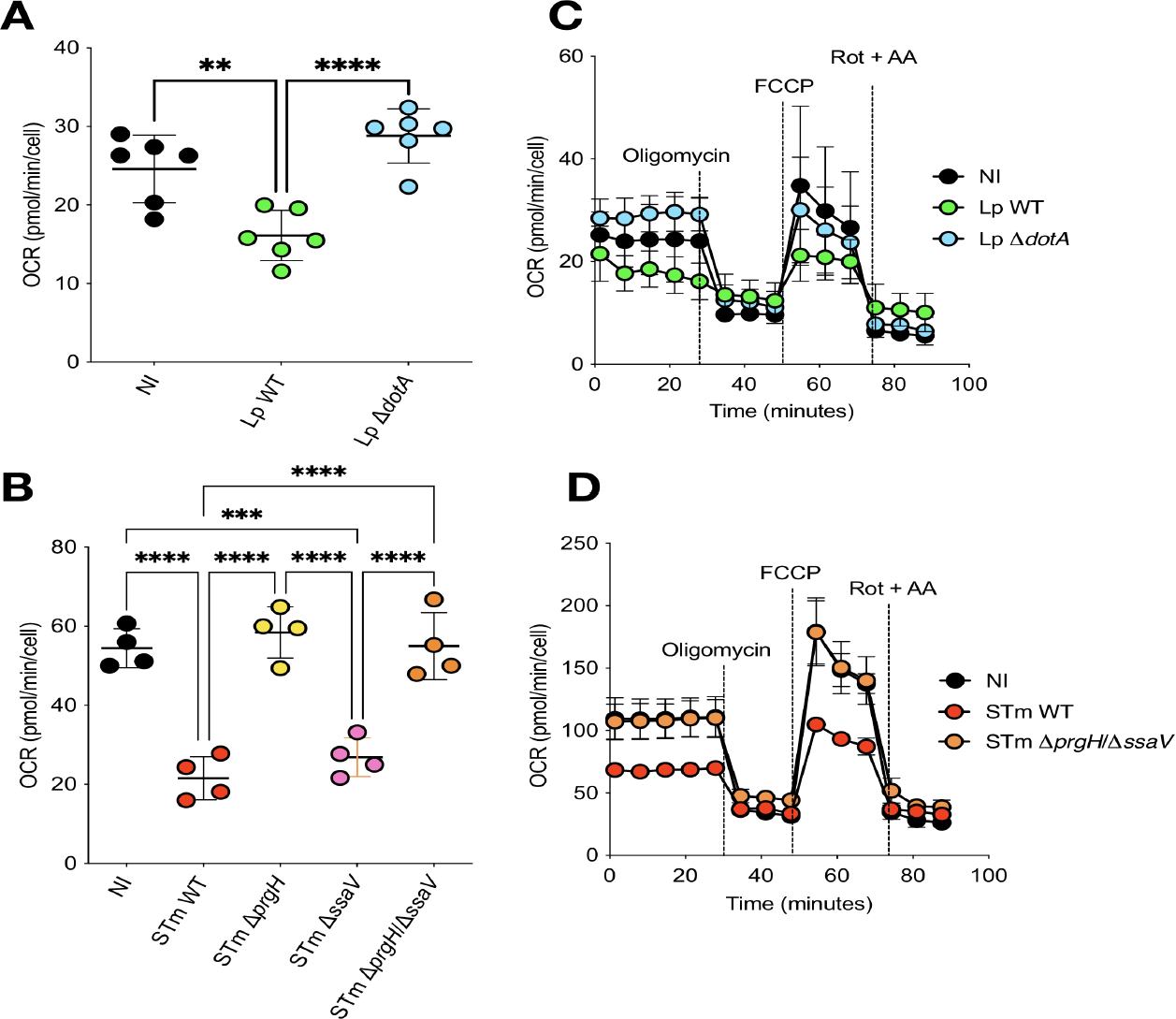
Secretion-dependent reduction of mitochondrial respiration in hMDMs at 5 hpi. (**A**) Basal oxygen consumption rate (OCR) in hMDMs infected for 5 h with *Legionella pneumophila* (*Lp*) WT or Δ*dotA* (T4SS-deficient), compared to non-infected controls (NI). (**B**) Basal OCR in hMDMs infected with *Salmonella enterica* serovar Typhimurium (*S*.Tm) WT or isogenic T3SS SPI-1 (Δ*prgH*), T3SS SPI-2 (Δ*ssaV*), or double T3SSs SPI-1/SPI-2 (Δ*prgH*/Δ*ssaV*) mutants. (**C**,**D**) Full Seahorse Mitostress profiles at 5 hpi for *Lp* (C) and *S*.Tm (D), with sequential additions (horizontal bars) of Oligomycin (ATP synthase inhibitor), FCCP (uncoupler), and Rotenone/Antimycin A (Rot+AA, Complex I/III inhibitors) to reveal ATP-linked respiration (difference between baseline and profile after adding Oligomycin), maximal respiration (difference between Oligomycin and profile after adding FCCP), spare capacity (difference between baseline and profile after adding FCCP), and non-mitochondrial respiration (profile after adding Rot+AA). Graphs show representative experiments from at least 3 independent experiments.

### Despite reduced mitochondrial respiration, infected macrophages preserve the mitochondrial membrane potential

To determine whether the reduction of OCR observed at 5 hpi is accompanied by the loss of the Δψm, we used Tetramethylrhodamine, methyl ester (TMRM), a dye for living cells that is retained in mitochondria proportionally to their Δψm. Thus, we quantified TMRM fluorescence in single cells as readout of the Δψm in hMDMs infected with *Lp*-WT or Δ*dotA* and with *S*.Tm-WT or Δ*prgH*/Δ*ssaV* (**Figure 2A**). Our analyses showed that the Δψm in *Lp*-WT- and *Lp*-Δ*dotA*-infected cells remained as high as in non-infected controls (**Figure 2B**), as previously shown ^19^. Likewise, *S*.Tm-WT- and *S*.Tm-Δ*prgH*/Δ*ssaV*-infected macrophages maintained the TMRM signal comparable to controls (**Figure 2B**). This aligns with previous reports showing T3SS-dependent metabolic reprogramming towards lower OxPhos levels and the preservation of the Δψm through the T3SS effector SipA in bone marrow-derived macrophages (BMDMs) ^20,16^. Our results extend now this observation also to human primary macrophages. Thus, despite the robust, secretion-dependent reduction in respiration, the Δψm is preserved at 5 hpi across infections with both bacterial species.

**Figure 2.**
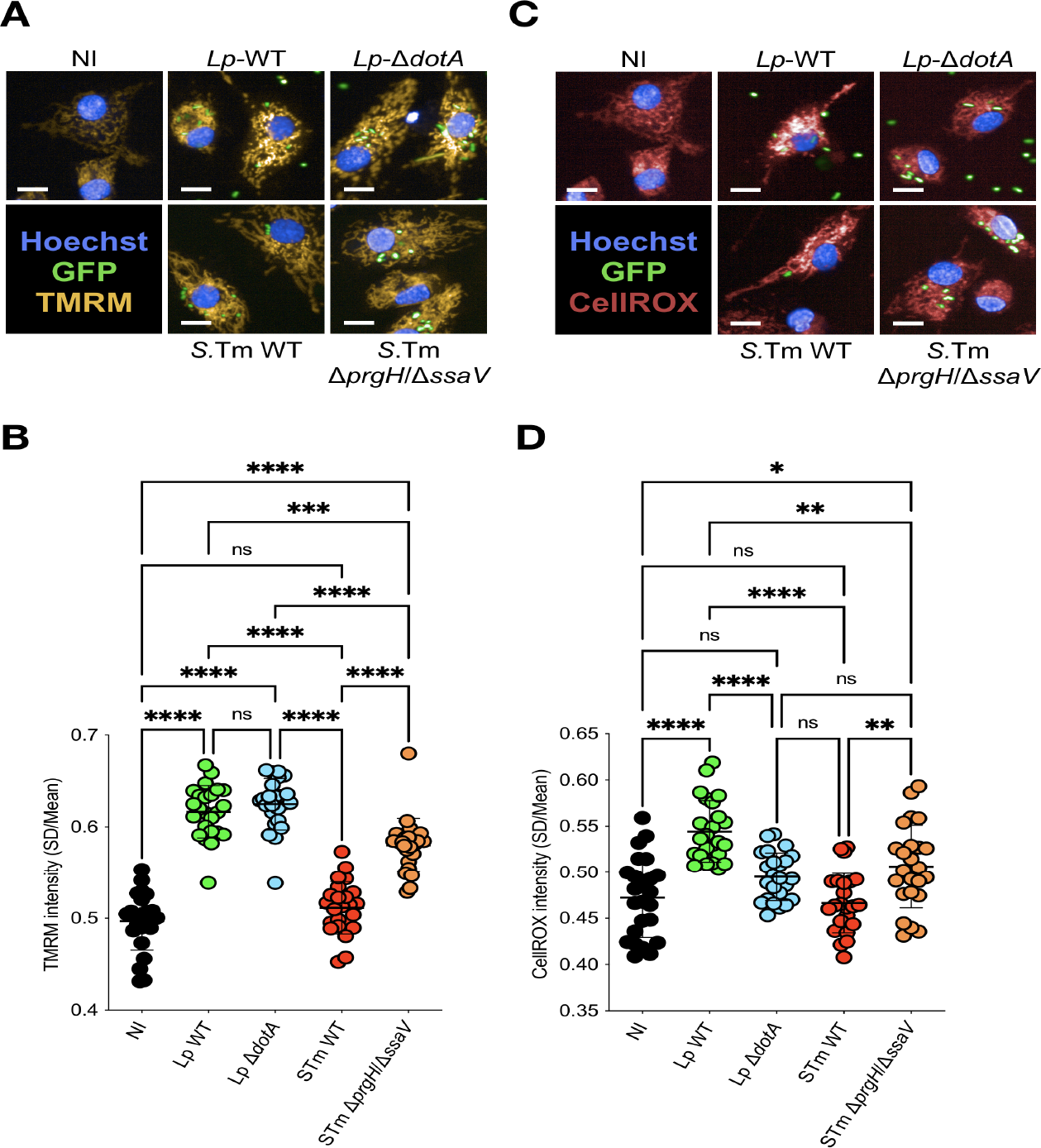
Despite reduced respiration, infected macrophages preserve Δψm while eliciting pathogen-specific ROS outputs. (**A**) Representative images of TMRM (Δψm) signals at 5 hpi in NI, *Lp*-WT, *Lp*-Δ*dotA, S*.Tm-WT, and *S*.Tm Δ*prgH*/Δ*ssaV* hMDMs. Bar = 10 µm. (**B**) Quantification of single-cell TMRM levels. Each dot is the mean value of all infected cells in a well. (**C**) Representative images of CellROX (mROS) signals at 5 hpi in NI, *Lp*-WT, *Lp*-Δ*dotA, S*.Tm-WT, and *S*.Tm Δ*prgH*/Δ*ssaV* hMDMs. Bar = 10 µm. (**D**) Quantification of single-cell mROS levels. Each dot is the mean value of all infected cells in a well. Plots reflect pooled mean values per well from single-cell measurements in a representative experiment from at least 3 independent experiments.

To investigate if the maintenance of the Δψm is accompanied by a production of mROS, we used CellROX, a dye for living cells that serves as readout of mROS accumulation in the mitochondria of human macrophages ^22^ (**Figure 2C**). Our measurements revealed pathogen-specific mROS patterns. While *Lp*-WT increased mROS production relative to non-infected and *Lp*-Δ*dotA* infected hMDMs, *S*.Tm-WT did not show a comparable increase, and hMDMs infected with the *S*.Tm-Δ*prgH*/Δ*ssaV* strain showed similar levels as non-infected cells (**Figure 2D**). Altogether, our results indicate that early during infection, virulence systems from both pathogens uncouple the respiratory flux from mitochondrial polarization, while eliciting distinct mROS outputs.

### Operational activity of the F_O_F_1_-ATPase differs between macrophages infected with Legionella pneumophila and with Salmonella Typhimurium

Previous work established that *Lp* preserves the Δψm in human macrophages by forcing the mitochondrial F_O_F_1_-ATPase to run in the ATP-hydrolyzing, reverse direction, a T4SS-dependent adaptation that compensates for the pathogen-driven drop in electron flow through the respiratory chain ^19^. In forward mode, protons flow through the F_O_ channel to drive ATP synthesis from ADP (**Figure 3A**). In reverse mode, F_1_ hydrolyses cytosolic ATP and F_O_ pumps protons out of the matrix to sustain the Δψm. Because the adenine-nucleotide translocator (ANT) exchanges ATP in the matrix for ADP in the cytoplasm, the reverse mode of the F_O_F_1_-ATPase is predicted to flip ANT towards importing ATP into the matrix (**Figure 3A**). ANT inhibition by bongkrekic acid (BA) therefore provides a functional readout of ANT coupling, whereas dicyclohexylcarbodiimide (DCCD) blocks proton translocation through the F_O_ channel, collapsing the Δψm specifically when the F_O_F_1_-ATPase is working in reverse mode (**Figure 3A**).

**Figure 3.**
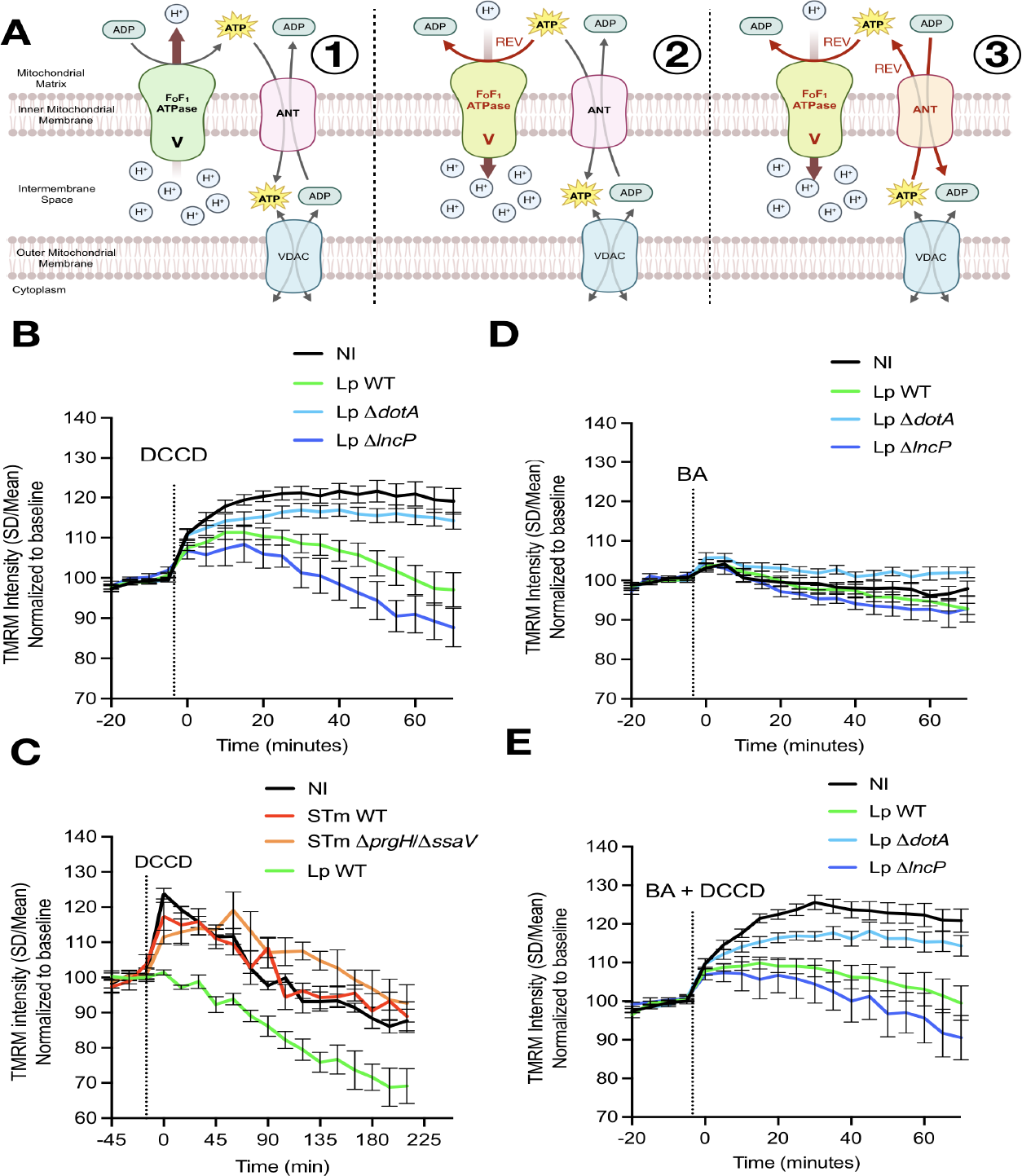
The operational mode of the F_O_F_1_-ATPase differs between *Lp* and *S*.Tm infection. (**A**) Schematic of forward (black arrows and text, FWD) vs reverse (red arrows and text, REV) operation of the F_O_F_1_-ATPase and ANT. In F_O_F_1_-ATPase_FWD_/ANT_FWD_ mode (1), protons re-enter via F_O_ to drive ATP synthesis and ANT transport ADP to the mitochondrial matrix and ATP to the cytoplasm via VDAC. In F_O_F_1_-ATPase_REV_/ANT_FWD_ mode (2), ATP hydrolysis by F_1_ pumps protons to the intermembrane space to sustain Δψm. ANT exchanges matrix ATP for cytosolic ADP. If ANT reverses in the F_O_F_1_-ATPase_REV_/ANT_REV_ mode (3), it can import ATP from the cytoplasm to fuel the reverse mode of F_O_F_1_-ATPase. (**B**) Δψm (TMRM) after addition of DCCD at 5 hpi in NI, *Lp*-WT, *Lp*-Δ*dotA*, and *Lp*-Δ*lncP*-infected hMDMs. (**C**) Same as B but during infection with *S*.Tm-WT or double mutant. (**D**) Same as B but adding BA. (**E**) Same as B but adding BA + DCCD combined. Graphs show representative experiments from at least 3 independent experiments.

To reveal whether the F_O_F_1_-ATPase and ANT worked in forward or reverse mode during infection, we monitored the Δψm in live hMDMs using TMRM and added DCCD, BA, or both. While a drop in Δψm upon DCCD addition would mean that the F_O_F_1_-ATPase worked in the reverse mode (hy-drolase activity), a drop in Δψm upon BA addition would mean that ANT worked in the reverse mode (importing ATP to the matrix). A bigger drop in Δψm upon simultaneous addition of DCCD and BA, compared to DCCD alone, would corroborate that the F_O_F_1_-ATPase and ANT both worked in reverse and are coupled to fuel the maintenance of Δψm with ATP imported from the cytoplasm of the cell. We also included in our assays the *Lp*-Δ*lncP* mutant because the T4SS effector LncP was reported to function as a mitochondrial carrier that is targeted to the inner mitochondrial membrane and transports ADP/ATP ^23^, which might influence ANT operations.

Addition of DCCD at 5 hpi during *Lp* infection elicited a rapid hyperpolarization in non-infected and in Δ*dotA*-infected macrophages, consistent with the forward-mode of the F_O_F_1_-ATPase now acting as a proton leak that is blocked by the inhibitor and protons now accumulate at the inter-membrane space. However, TMRM dropped by 25 % within 60 min in WT-infected macrophages, demonstrating that the Δψm depends on the reverse mode of the F_O_F_1_-ATPase during *Lp*-WT infection (**Figure 3B**), as previously shown ^19^. We next investigated the operational mode of the F_O_F_1_-ATPase during *S*.Tm infection using DCCD (**Figure 3C**). *S*.Tm-WT and the T3SS SPI-1/SPI-2 double mutant (Δ*prgH*/Δ*ssaV*) behaved like non-infected cells, indicating that, unlike *Lp, S*.Tm preserves the Δψm without a major contribution from the reverse activity of the F_O_F_1_-ATPase.

To know if the reversal of the F_O_F_1_-ATPase during infection by *Lp* was accomapined by a reversal of ANT, we added BA alone at 5 hpi, which induced only a modest, and similar decline in TMRM in non-infected, *Lp*-WT- and *Lp*-Δ*dotA*-infected macrophages (**Figure 3D**), but caused a larger drop in the Δ*lncP* mutant, suggesting that ANT activity is not a dominant determinant of the Δψm in non-infected, WT- or Δ*dotA*-infected cells but becomes partially engaged when LncP is absent. When BA and DCCD were added together (**Figure 3E**), the TMRM profile of *Lp*-WT infected cells was indistinguishable from the trace obtained with DCCD alone (**Figure 3B**), consistent with ANT remaining in forward mode at 5 hpi. Non-infected and Δ*dotA* infected cells again showed the mild hyperpolarization characteristics for the forward mode of the F_O_F_1_-ATPase. The lack of an additional drop of the Δψm corroborated that, at 5 hpi, ANT is still exchanging matrix ATP for cytosolic ADP (forward mode of ANT). Thus, the reverse mode of function of the F_O_F_1_-ATPase during *Lp*-WT infection alone does not depolarize the membrane enough to reach the ANT reversal potential. Previous thermodynamic analyses placed the reversal potential of the F_O_F_1_-ATPase at more negative values than that of ANT ^24,25^, meaning that the F_O_F_1_-ATPase flips first, and its proton-extruding activity counteracts further depolarization that would otherwise force ANT to import ATP. According to this framework, a deeper or more sustained depolarizing action, such as a stronger reverse ATPase activity or prolonged infection, would be required before BA synergizes with DCCD dropping the Δψm further. Thus, we cannot exclude ANT reversal at later times or under stronger reverse of the F_O_F_1_-ATPase. Interestingly, the *Lp*-Δ*lncP* mutant strain showed an increased drop of the Δψm compared to WT upon adding DCCD or DCCD + BA, which suggests that LncP shapes nucleotide exchange in a way that the reverse mode of the F_O_F_1_-ATPase is attenuated, such as increasing the flux of ADP from the cytoplasm to the mitochondrial matrix.

Taken together, our results indicate that *Lp* induces the reverse operation of the F_O_F_1_-ATPase early after infection but keeps ANT in forward mode, whereas *S*.Tm preserves mitochondrial polarization without substantial reverse F_O_F_1_-ATPase activity. These results support a pathogen-specific modulation of the host cell bioenergetics where the working of the F_O_F_1_-ATPase and the nucleotide exchange machinery sustain the Δψm while minimizing deleterious cytosolic ATP consumption.

### Complex I-driven maintenance of the Δψm differentiates secretion-competent, virulent from secretion-deficient bacteria

To understand which complexes of the respiratory-chain sustain the the Δψm during infection, we monitored single-cell TMRM fluorescence in hMDMs at 5 hpi while inhibiting individual ETC complexes (**Figure 4A**). Rotenone (inhibitor of Complex I), thenoyltrifluoroacetone (TTFA, inhibitor of Complex II), antimycin A (inhibitor of Complex III) or sodium cyanide (NaCN, inhibitor of Complex IV) were added, and the levels of TMRM intensity of every single, infected macrophage were recorded for 200 min. In parallel, CellROX was tracked after addition of Rotenone to quantify mROS production by Complex I as done previously ^22^.

**Figure 4.**
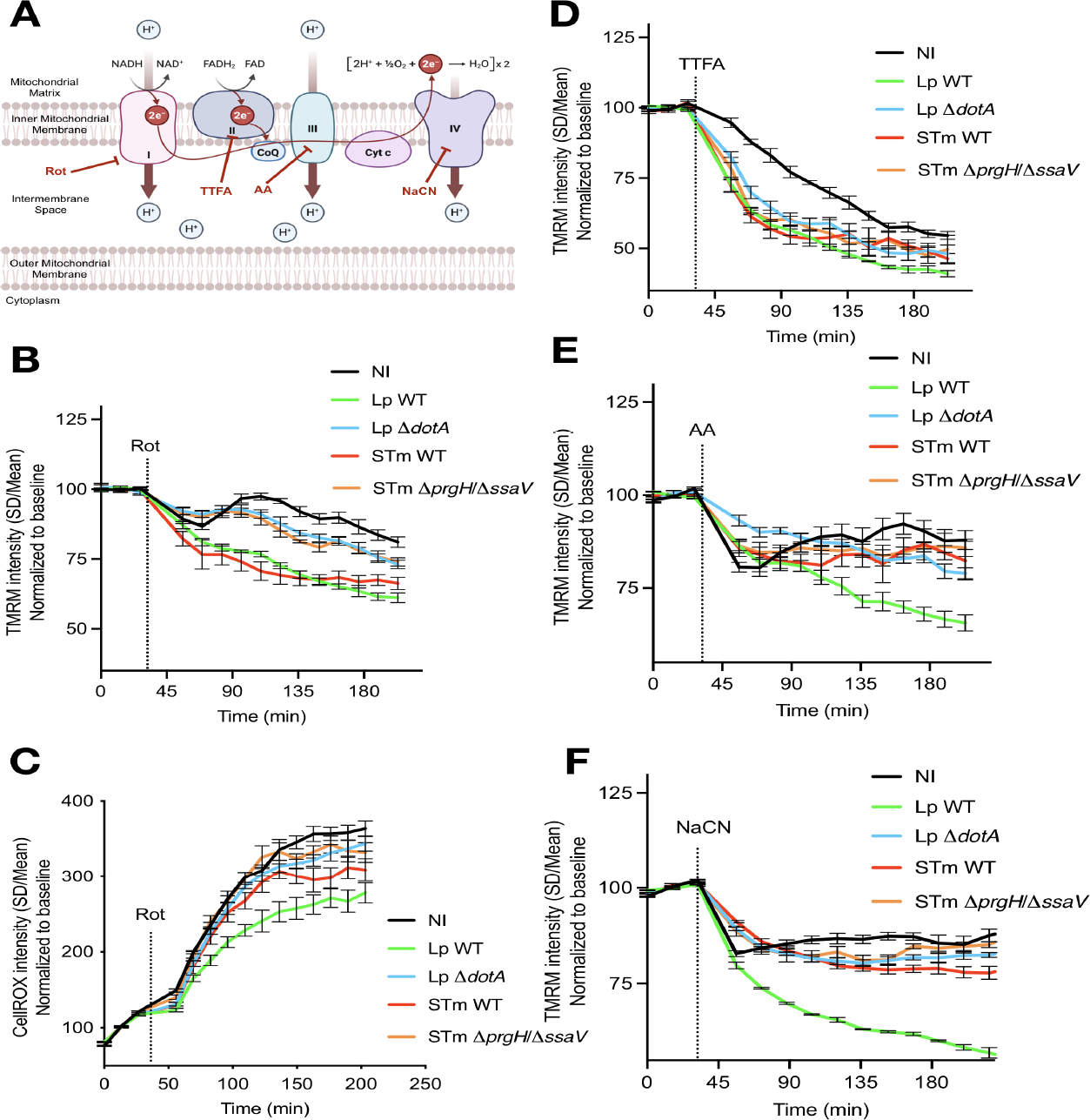
Complex I uniquely marks virulent infection while Complex II is recruited by infection per se and Complexes III–IV are universally required. (**A**) Schematic of the ETC (Complexes I-IV) and the F_O_F_1_-ATPase indicating inhibitor targets: Rotenone (Rot, Complex I), thenoyltrifluoroacetone (TTFA, Complex II), antimycin A (AA, Complex III), and sodium cyanide (NaCN, Complex IV). (**B**) Δψm (TMRM) after addition of Rotenone at 5 hpi in NI, *Lp*-WT, *Lp*-Δ*dotA, S*.Tm-WT and *S*.Tm-Δ*prgH*/Δ*ssaV* double mutant. (**C**) Same as B but measuring mROS. (**D**) Same as B but adding TTFA to assess Complex II contribution. (**E**) Same as B but adding Antimycin A to assess Complex III contribution. (**F**) Same as B but adding NaCN to assess Complex IV contribution. Graphs show representative experiments from at least 3 independent experiments.

Complex I inhibition revealed a unique, virulence-dependent activity. Rotenone triggered a pronounced drop of 35 % in TMRM levels in *Lp*-WT and in *S*.Tm-WT, whereas non-infected cells depolarized by only 15 %, and secretion-deficient mutants (Δ*dotA*, Δ*prgH*/ΔssaV) showed minimal additional loss (**Figure 4B**). The same Rotenone pulse markedly increased mROS production in infected macrophages, but less in *Lp*-WT than in the other infection conditions (**Figure 4C**), indicating that the Complex I-derived electron flow is the major proton source that sustains the Δψm during WT infection and suggesting that the already increased basal mROS production in *Lp*-WT-infected macrophages (**Figure 2D**) might have exhausted the host cells to increase mROS further upon Rotenone challenge.

Blocking Complex II with TTFA decreased the Δψm in all infection conditions, but had little effect on non-infected cells (**Figure 4D**), suggesting that Complex II supports the Δψm only when bacterial infection, with virulent or avirulent strains, perturbs metabolic fluxes. Inhibiting Com-plex III (antimycin A) or Complex IV (NaCN) collapsed the Δψm across all groups (**Figure 4E** and **4F**), consistent with their essential proton-pumping roles irrespective of the infection state but showing a bigger effect on *Lp*-WT-infected macrophages.

These patterns indicate that the presence of secretion systems channels the flow of electrons in the mitochondria of host cells through Complex I to maintain the Δψm and probably to also generate mROS. The strong mROS burst after Rotenone addition supports Complex I engagement in all infection conditions but specially in *Lp*-WT-infected cells. Secretion-deficient mutants lack the CI-dependent signature and show a greater relative impact of CII inhibition, consistent with infection-driven recruitment of CII rather than virulence per se. The uniform sensitivity of the Δψm to Antimycin A and NaCN treatments, inhibitors of Complex III and Complex IV, respectively, underscores that Complex III and Complex IV remain essential for keeping the Δψm in all infection conditions. Thus, the responses to specific com-plex inhibitors show that Complex I is a bioenergetic checkpoint that separates virulent intracellular bacteria from their secretion-defective counterparts, whereas downstream proton pumps (Complex III and Complex IV) remain universally required.

The Complex I signature we observed here might be linked to a broader innate-immune metabolic network described previously ^26^. Recognition of living, Gram-negative bacteria by murine and human macrophages transiently destabilizes Complex I-containing super-complexes, lowers Complex I activity and, in compensation, boosts electron flux through Complex II ^26^. Our data extend that framework by showing that, in primary human macrophages, virulent *Lp* and *S*.Tm actively utilize Complex I-dependent proton pumping to preserve the Δψm, whereas secretion-deficient mutants or avirulent bacteria use Complex II in a configuration reminiscent of the viability-sensing state described in murine macrophages ^26^. Thus, the engagement of Complex I emerges as both a marker of pathogen virulence and of a potential metabolic handle for modulating the host defense.

Our results support a model (**Figure 5**) in which *L. pneumophila* reprograms human macrophage bioenergetics along two coordinated axes. First, the T4SS effector MitF dampens OxPhos early after entry via fragmenting mitochondria networks ^18^. Second, the effector LpSpl participates to drive the F_O_F_1_-ATPase into its ATP-hydrolyzing reverse mode ^19^, pumping protons out of the matrix to preserve the Δψm without triggering cell death (**Figure 5A**). Because this reverse activity effectively maintains the Δψm, the adenine-nucleotide translocator (ANT) continues to operate in its forward mode, importing ADP from the cytoplasm into the ma-trix, exporting ATP from the matrix to the cytoplasm, and preventing cytosolic ATP depletion, a configuration likely stabilized by the T4SS effector LncP. All ETC complexes (I-IV) help to sustain the Δψm under these conditions, but Complex I is not required when secretion systems are absent.

**Figure 5.**
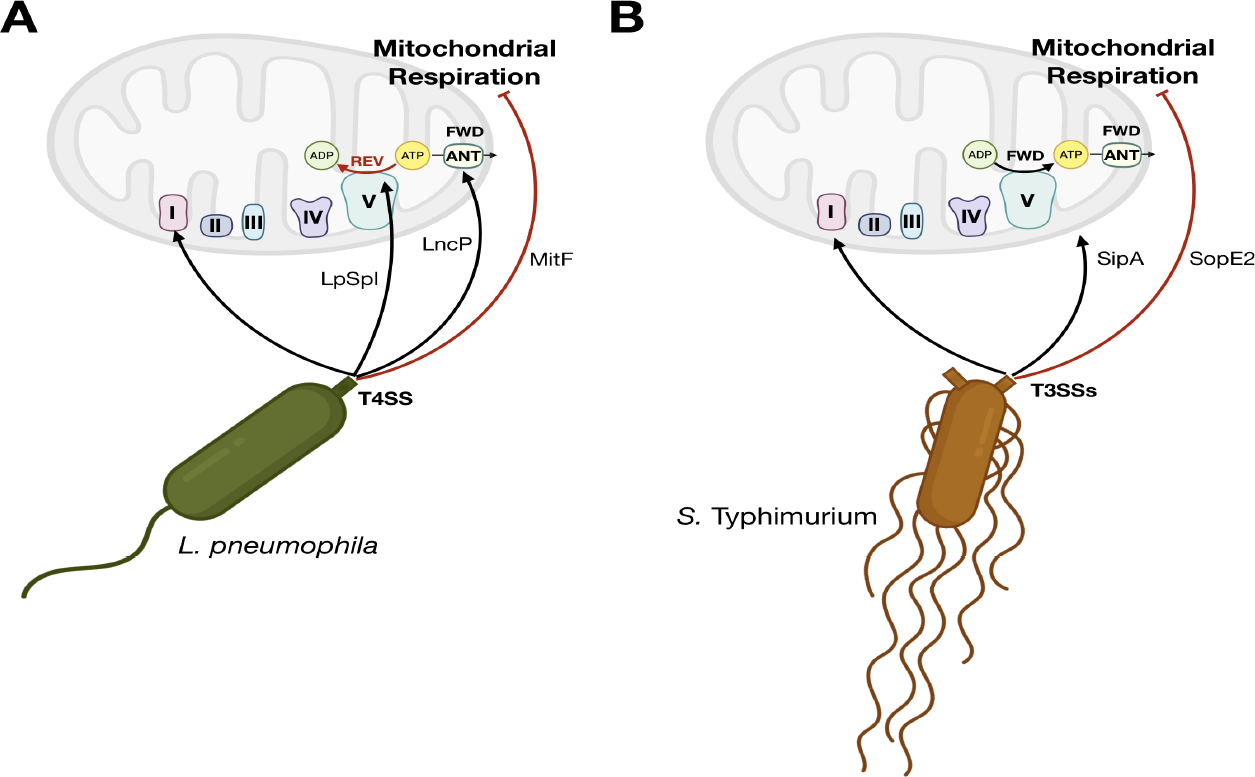
Working model for pathogen-specific ETC modulations that preserve the Δψm in hMDMs. (**A**) *L. pneumophila*: T4SS effectors dampen OxPhos early during infection and preserve the Δψm by engaging the reverse mode of the F_O_F_1_-ATPase. The Δψm remains high enough that ANT stays in forward mode, limiting cytosolic ATP drain. All ETC complexes (I-IV) contribute to maintaining the Δψm. Complex I is uniquely essential during WT infection. (**B**) *S*. Typhimurium: T3SS SPI-1/2 effectors reduce respiration and preserve the Δψm without inducing the reverse mode of the F_O_F_1_-ATPase. Virulent bacteria rely on Complex I-driven electron entry with obligatory Complex III/IV proton pumping, while Complex II contributes in all infection conditions. Secretion-defective mutants fail to establish the Complex I-dependent configuration.

*Salmonella* Typhimurium follows a different path (**Figure 5B**). Its SPI-1 T3SS effector SopE2 reduces mitochondrial respiration and the SPI-2 effector SipA maintains the Δψm via promoting the elongation of mitochondria ^20,16^, without flipping the F_O_F_1_-ATPase into reverse mode. Virulent bacteria channel the electron flow predominantly through all ETC complexes, but secretion-defective (Δ*prgH*/Δ*ssaV*) mutants fail to engage the here described Complex I-driven circuit, and they thus resemble the non-virulent *L. pneumophila* Δ*dotA* mutant phenotype. Together, these findings reveal a pathogen-specific hierarchy of ETC modulation. While Complex I activity delineates infection by virulent, intracellular bacteria, the ability to reverse the ATP-synthase activity to sustain Δψm distinguishes the ETC functioning patterns of macrophages infected with *L. pneumophila* from those infected by *S*. Typhimurium.

## Acknowledgements

We acknowledge all C.B’s lab members for fruitful discussions. We thank Nathalie Aulner, Anne Danckaert, Nassim Mahtal and the Photonic BioImaging (PBI) UTechS at Institut Pasteur for their support. We are grateful to the healthy volunteers for their participation in the study. We acknowledge the Collections for Human Health of the Institut Pasteur biobank (CHIP) of the Biological Resources Center of the In-stitut Pasteur (CRBIP) for the distribution of the blood samples. This research was funded by the Institut Pasteur, the Agence National de Recherche (ANR-21-CE15-0038-01 to P.E and ANR-10-LABX-62-717 IBEID to C.B), the ‘Fondation pour la Recherche Médicale’ (grant EQU201903007847 to C.B), the ‘Programmes Transversaux de Recherche’ (PTR-651) from Institut Pasteur to P.E and the Région Ile-de-France (program DIM1Health) to PBI (part of FranceBioImaging, ANR-10-INSB-04–01). Figures have elements from Biorender.

## Methods

### Bacterial strains and plasmids

All bacterial strains used in this study expressed GFP from plasmid pNT28 (*Legionella pneumophila*) or pFPV25.1 (*Salmonella enterica* serovar Typhimurium) and carried the corresponding antibiotic resistance cassette (Table S1). *L. pneumophila* strain Paris wild type and isogenic mutants lacking either the type IV secretion system (Δ*dotA*) or the effector LncP (Δ*lncP*) were cultured under identical conditions. Bacteria were maintained on ACES-buffered charcoal-yeast extract (BCYE) agar supplemented with chloramphenicol (10 µg/mL; Sigma) at 37°C for 3 days. For liquid culture, colonies were inoculated into 10 mL of Buffered Yeast Extract (BYE) medium containing chloramphenicol (10 µg/mL) at an initial OD_600_ of 0.1 and incubated overnight at 37°C with shaking (200 rpm). Cultures were harvested at late exponential phase (OD_600_ 4.2) and diluted in cell culture medium to reach the desired multiplicity of infection (MOI = 10).

*Salmonella enterica* serovar Typhimurium strains were derived from the parental ATCC 14028 background and included wild type, Δ*prgH*, Δ*ssaV*, and the double mutant Δ*prgH*/Δ*ssaV* (listed in Table S1). Bacteria were routinely grown in Lysogeny Broth (LB) supplemented with ampicillin (100 µg/mL).

### Host cells

hMDMs isolation and differentiation were performed as previously described ^30^. Blood from healthy donors was provided by the Clinical Investigation Center INVOLvE (Investigation and Volunteers for Human Health) of the Institut Pasteur. All participants received oral and written information about the research and gave written informed consent in the frame of the healthy volunteers COSIPOP cohort after approval of the CPP Est II Ethics Committee (2023, February 20th). Peripheral blood mononuclear cells (PBMCs) were isolated by Ficoll density gradient centrifugation (Lymphocyte Separation Medium, Eurobio Scientific) at room temperature. Monocytes were positively sorted using CD14 magnetic beads (CD14 MicroBeads, Miltenyi Biotec). Following, monocytes were differentiated into hMDMs by culturing them in X-VIVO15 medium without Phenol Red (Lonza) in Nunc Up-Cell plates (Thermo Scientifics) and in the presence of recombinant human M-CSF (rh-MCSF, Miltenyi Biotec) at a concentration of 25 ng/ml for 6 days at 37°C with 5 % CO_2_ in a humidified incubator. The day before infection, cells were detached from UpCell plates and plated in 384-well plates (Greiner Bio-One) at a cell density of 10,000 cells per well in X-VIVO15.

### Infections of host cells with *L. pneumophila* and *S*. Typhimurium

hMDMs were infected with either *Lp* or *S*.Tm under conditions designed to promote synchronized uptake and restrict extracellular replication. For *Lp*, one the day prior to the infection, 10 mL BYE liquid medium with 10 µg/mL of chloramphenicol were inoculated with the bacteria at an optical density of OD_600_ = 0.1. The bacterial culture was allowed to grow overnight with shaking at 37°C and 200 rpm until OD_600_ reached approximately 4.2. Then, the bacterial culture was diluted in the final cell culture medium to achieve a multiplicity of infection (MOI) of 10.

For *S*.Tm, a single colony from an LB agar plate supplemented with 100 µg/mL ampicillin was used to inoculate LB broth containing 100 µg/mL ampicillin and grown overnight at 37°C with shaking; on the day of infection, 150 µL of this culture was diluted 1:33 into 5 mL of fresh LB medium and incubated with shaking for 3 h to reach early stationary phase, after which bacteria were harvested, washed in serum-free medium, resuspended in X-VIVO15, and used to infect hMDMs at an MOI of 15.

Infections were synchronized for both pathogens by centrifugation of plates at 200 × g for 5 min at room temperature, immediate transfer to a 37°C water bath for 5 min to facilitate phagocytosis, and a subsequent 25-min incubation at 37°C in a humidified atmosphere with 5 % CO_2_; cells were then washed extensively with PBS (3-5 times) to remove non-internalized bacteria, treated with medium containing 100 µg/mL gentamicin for 20 min to eliminate residual extracellular bacteria, and subsequently maintained in culture medium containing 10 µg/mL gentamicin for the duration of the experiment to prevent extracellular replication.

### Electron Transport Chain drugs

To assess the contributions of mitochondrial ETC complexes and accessory proteins to Δψm during infection, hMDMs were treated at 5 h post-infection with the F_O_F_1_-ATPase inhibitor DCCD (100 µM, Sigma), the ANT inhibitor Bongkrekic Acid (BA, 25 µM, Sigma), the Complex I inhibitor Rotenone (10 µM, Sigma), the Complex II inhibitor TTFA (1 mM, Sigma), the Complex III inhibitor Antimycin A (10 µM, Sigma) or the Complex IV inhibitor NaCN (1 mM, Sigma).

### Staining of living cells and automated time-lapse confocal imaging acquisition

For automated time-lapse confocal imaging experiments, we performed infection assays in 384-well microplates as previously described ^30^. Each experiment was performed using four technical replicates and repeated to obtain at least three biological replicates. To track individual cells, nuclei and cytoplasm were stained one hour before infection with Hoechst H33342 (200 ng/mL, Life Technologies) and CellTracker Blue (25 µM, Invitrogen), respectively. To measure mitochondrial membrane potential and ROS, TMRM (25 nM, non-quench mode, Life Technologies) and CellROX Deep Red (5 µM, Life Technologies) were added to the culture medium and maintained dur-ing the assays. Image acquisition was performed using an automated microlens-enhanced spinning disc confocal microscope (Opera Phenix High Content Screening System, PerkinElmer) equipped with 40x and 60x water objectives. Excitation lasers operated at 405, 488, 561, and 640 nm, with emission filters set at 450, 540, 600, and 690 nm. To capture the dynamics of mitochondrial parameters, images of multiple fields (ranging from 10 to 25) were acquired in time-lapse experiments using an incubation chamber maintained at 37°C with 5 % CO_2_.

**Table 1.** List of bacterial strains used in this study.

| Strain (Plasmid) | Relevant genotype/description | Antibiotic Selection | Ref. |
| --- | --- | --- | --- |
| *Lp*-WT-GFP (pNT-28) | Wild-type *Legionella pneumophila* strain Paris | Chloramphenicol | 27,28 |
| *Lp*-Δ*dotA*-GFP (pNT-28) | T4SS-defective mutant of *Legionella pneumophila* Paris | Chloramphenicol | 22 |
| *Lp*-Δ*lncP*-GFP (pNT-28) | LncP-defective mutant of *Legionella pneumophila* Paris | Chloramphenicol | 19 |
| *S*.Tm-WT-GFP (SV9760/pFPV25.1) | Wild-type *Salmonella enterica* ser. Typhimurium 14028 | Ampicillin | 29 |
| *S*.Tm-Δ*prgH*-GFP (SV5379/pFPV25.1) | T3SS SPI-1-defective *S*.Tm 14028 | Ampicillin | 29 |
| *S*.Tm-Δ*ssaV*-GFP (SV5136/pFPV25.1) | T3SS SPI-2-defective *S*.Tm 14028 | Ampicillin | 29 |
| *S*.Tm-Δ*prgH*Δ*ssaV*-GFP (SV6747/pFPV25.1) | T3SS SPI-1/2-defective *S*.Tm 14028 (double mutant) | Ampicillin | 21 |

### Image Analyses

Quantitative image analysis was performed using Harmony software v4.9 (PerkinElmer) complemented by custom-built scripts (shared upon request). Nuclei were segmented based on Hoechst fluorescence, while cytoplasmic regions were defined using either CellTracker Blue, TMRM, or CellROX background signals depending on the analysis. Intracellular bacteria were detected by their constitutive GFP signal. Mitochondrial membrane potential (Δψm) was quantified from TMRM fluorescence (red channel) and intracellular ROS levels were measured from CellROX fluorescence (far-red channel), both with normalized per-cell values calculated as the ratio of standard deviation to mean intensity (SD/mean).

### Real-time extracellular flux assays (Seahorse)

OCR of macrophages were determined using an XF96 extracellular flux analyzer (Seahorse Bioscience, Agilent). hMDMs (6 × 10^5^ cells/well) were plated in XF96-cell culture plates containing XVIVO15, infected with *Lp* or *S*.Tm and incubated for 5 h at 37°C and 5% CO_2_. One hour before analysis, the culture medium was removed from the cells, and the cells were incubated at 37°C and 0 % CO_2_ in XF assay RPMI medium containing 10 % (v/v) FBS supplemented with 10 mM glucose, 1 mM pyruvate and 2 mM glutamine. OCR values were taken before and after the sequential addition of Oligomycin (2.5 µM), FCCP (2 µM) and Rotenone + Antimycin A (both 0.5µM))XF Mito Stress Kit, Seahorse Bioscience, Agilent). Basal OCR values were obtained by subtracting the non-mitochondrial OCR values, respectively, and after normalizing to the cell number.

### Generative AI statement

During the preparation of this work, authors used ChatGPT 4.5 (OpenAI) and Gemini 2.5pro (Google) Large Language Models (LLMs) in order to correct grammar and flow of some parts of the text. After using these tools, authors reviewed and edited the content as needed and take full responsibility for the content of the publication.

